# Genomic and evolutionary factors influencing the prediction accuracy of optimal growth temperature in prokaryotes

**DOI:** 10.1101/2025.05.30.656958

**Authors:** Seiji Toki, Motomu Matsui, Kento Tominaga, Takao K. Suzuki, Takashi Tsuchimatsu, Wataru Iwasaki

## Abstract

Bacteria and archaea have evolved diverse genomic adaptations to thrive across various temperatures. These adaptations include genome sequence optimizations, such as increased GC content in rRNA and tRNA, shifts in codon and amino acids usage, and the acquisition of functional genes conferring adaptation for specific temperatures. Since the experimental determination of optimal growth temperatures (OGT) is only possible for cultured species, predicting OGT from genomic information has become increasingly important given the exponential increase in genomic data. Although previous studies developed prediction models integrating multiple features based on genome composition using machine learning, the accuracy was variable depending on the target species, with models performing well for thermophiles but less accurately for psychrophiles. In this study, we curated the OGT and genomic data of 2,869 bacterial species to develop a novel prediction model incorporating features reflecting genomic adaptation toward lower temperatures. We found that species with rapid OGT shifts from their ancestors, including psychrophiles, showed less accuracy in genome composition-based models. Incorporating the gene presence/absence information associated with the rapid changes in OGT improved the prediction accuracy for psychrophiles. We also observed that OGT in archaea is phylogenetically more conserved than in bacteria, which may lead to the long-term optimization of the genome composition and explain high predictability of OGT in archaea. These findings highlight the importance of integrating long- and short-term evolutionary adaptations for phenotype prediction models.

## Importance

Prediction of optimal growth temperature (OGT) from genomic data allows for characterizing uncultivated microbes. This study developed a novel OGT prediction model by curating OGT and genomic data of 2,869 bacterial species, demonstrating that the prediction accuracy was improved by incorporating not only genome composition features reflecting long-term temperature adaptation but also the presence/absence information of genes conferring short-term adaptation, particularly for species undergoing rapid OGT shifts. Our study provides a framework for improving phenotype prediction models by integrating long- and short-term evolutionary factors, which may also apply to other microbial physiological phenotypes.

## Background

Bacteria and archaea have adapted to a wide range of temperatures from below 0 °C to over 100 °C^1, 2^, and their genomes have undergone diverse evolutionary modifications in response to their habitat temperatures. In general, bacterial thermal adaptation strategies can be classified into two major types: optimization of genome sequences^3^ and the gain of genes that confer specific functions^4^. Since genome sequences fundamentally determine the stability of nucleic acid secondary structures and the folding of the encoding proteomes, they should be optimized to specific habitat temperatures. For example, the GC content of rRNA and tRNA is elevated and the ratio of charged to polar amino acids increases in thermophiles^5–7^. In addition, synonymous codon usage for arginine varies in response to the species’ habitat^8^. The gain of genes also facilitates bacterial adaptation to specific temperatures. Some thermophiles possess reverse gyrases that introduce positive supercoil and stabilize DNA under high temperatures^9^. Similarly, psychrophiles often possess chaperones that facilitate proper RNA folding in cold environments and antifreeze proteins that inhibit the formation of ice crystals inside the cells^1^.

Although growth temperature is one of the most fundamental parameters of prokaryotic physiology, the experimental determination of prokaryotes’ optimal growth temperature (OGT) is laborious and applicable only to cultured species^10^. Therefore, predicting OGT from genomic information has become increasingly important given the exponential increase in genomic data from cultivated and uncultivated prokaryotes in recent years^11, 12^. For the prediction of OGT, the information on genome composition (i.e., base composition, codon usages, and amino acids composition) has been widely adopted because it does not require functional gene annotation and is applicable across a broad range of taxa. As a simple and reliable indicator of OGT, a previous study demonstrated that the fraction of seven amino acids—isoleucine, valine, tyrosine, tryptophan, arginine, glutamic acid, and leucine (abbreviated as IVYWREL)—shows a strong correlation with OGT^13^. Recently, machine learning has been incorporated to improve the prediction, integrating many genome-derived features, such as tRNA and rRNA base composition, codon usage, and amino acid composition, leading to the development of highly accurate prediction models^12, 14^. These studies demonstrate that OGT can be generally predictable based on genome composition.

Despite such methodological advancement in predicting OGT, its accuracy based on genome composition was variable depending on the target. While the model showed an improved fit for thermophiles, an accurate prediction of OGT for psychrophiles based on genomic data poses challenges, possibly because of the underrepresentation of species and/or features associated with the adaptation to low temperatures^12, 15^. Furthermore, a previous study reported higher correlations between genomic features and OGT in archaea than in bacteria, resulting in higher predictability of OGT in archaea^12^. However, the reasons for the differences in prediction accuracy between targets are largely unexplored.

In this study, we aimed to clarify the reasons for the difficulty in the OGT prediction from prokaryotic genomes, especially in psychrophiles, and provide a framework for improving accuracy. Using the curated OGT and genomic data of 2,869 bacterial species, we developed a novel prediction model incorporating features reflecting the genomic adaptation toward lower temperatures. We found that the model based on genome composition poses a challenge in predicting the OGT of species whose OGT has rapidly changed from their ancestors, including psychrophiles, and that the prediction accuracy of OGT in psychrophiles was improved by incorporating the gene presence/absence information associated with the rapid changes in OGT. Finally, we discovered that OGT in archaea is phylogenetically more conserved than in bacteria, which may lead to the long-term optimization of the genome composition and explain why the predictability of OGT is high in archaea. These results highlight the importance of considering traits’ long- and short-term evolution for developing highly accurate phenotype prediction models.

## Results

### Constructing an OGT prediction model based on genome composition

We first curated the OGT data and genomes of corresponding species from public databases. After merging OGT from three databases^16–18^ and the corresponding genomes of GTDB^19–22^, OGT and genome of 2,869 bacterial and 262 archaeal species were obtained (Figure 1a,b). In this study, we defined thermophiles as species with OGT ≤ 60 °C and psychrophiles as species with OGT < 20 °C according to the previous studies^23, 24^. Because OGT was most frequent from 20 to 40 °C (2,278 species) as previously reported^14^, the model may overfit this temperature range and underestimate psychrophile features. We downsampled the data (1,631 species) to verify this hypothesis by selecting one species per genus whose OGT is 20-40 °C.

**Figure 1.**
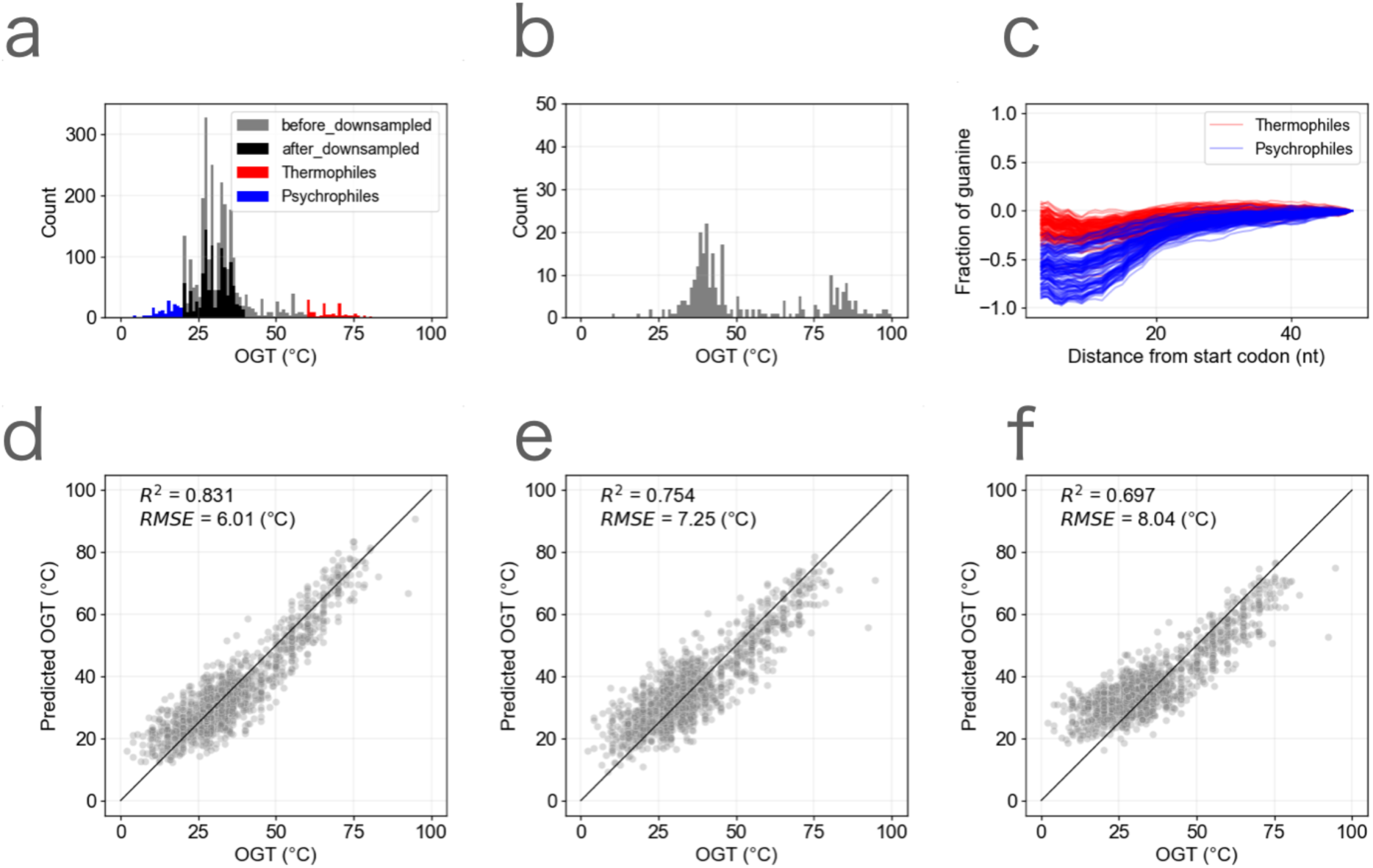
Constructing an OGT prediction model based on genome composition. **a** The distribution of OGT of bacteria (2,869 species). Red and blue indicate thermophiles (OGT ≥ 60 C°) and phychrophiles (OGT < 20 C°), respectively. Black indicates the distribution after downsampling mesophiles (20 C° ≤ OGT < 40 C°). **b** The distribution of OGT of archaea (262 species). **c** The fraction of guanine in the downstream region of the start codon. The fraction was normalized at 50 nt downstream of the start codon by taking the logarithm to base 2. The red and blue lines indicate thermophiles and psychrophiles, respectively. **d-f** The comparison of the prediction models based on genome composition. Each dot represents one bacterial species. The predictions based on genome composition using Lasso regression **(d**) and SVR (**e**) are shown. (**f)** The prediction using the model of Li et al., 2019 (replicated in this study).

Next, we calculated genomic features used for OGT prediction. We first computed features similar to those used in previous studies^12, 14^, including tRNA base composition, rRNA base composition, amino acid composition, and codon composition (Supplementary Table 1). While the GC content of tRNA and rRNA is known to correlate with temperature^3, 5, 6^, we also calculated the minimum free energy (MFE) of tRNA and rRNA that showed a stronger correlation with OGT than GC content (Supplementary Figure 1), suggesting that the MFE is the feature that better reflects the secondary structure of RNA. To improve the prediction of psychrophiles, we also calculated the features around the start codon, because a stronger selection pressure would be exerted around the start codon of the mRNA sequence at lower temperatures, given the inhibition of the initiation of translation by the secondary structures^25, 26^. Consistently, we found that the frequency of guanine significantly decreases in psychrophiles compared with thermophiles (Figure 1c), incorporating the frequency of guanine and amino acids around the start codon for the prediction (21 features, Supplementary Figure 2).

We developed the OGT prediction model using the features described above. To verify if the model can predict the OGT of clades that are not used for the training dataset, a leave-one- phylum-out cross-validation was performed, in which one phylum was used as the test data and the rest were used as the training data. We found that Lasso regression showed higher accuracy (*R*^2^ = 0.831, *RMSE* = 6.01 °C) than support vector regression (SVR) (*R*^2^ = 0.754, *RMSE* = 7.25 °C) (Figure 1d,e). Although the most accurate prediction model to date used SVR based on the amino acid 2-mer composition^14^, the model reproduced was also less accurate compared with our models (*R*^2^=0.697, *RMSE*=8.04 °C, Figure 1f).

Despite the improvement in overall accuracy, the accurate prediction of OGTs for psychrophiles remained challenging (*mean absolute error* = 7.1°C) compared with the accuracy of other temperature ranges (*mean absolute error* = 4.4 °C). The prediction of OGTs for psychrophiles became worse without downsampling the data (*mean absolute error* = 8.0 °C), without features around the start codon (*mean absolute error* = 7.7 °C), or without both procedures (*mean absolute error* = 8.8 °C). These results indicate that balancing the distribution of OGT in the training dataset and incorporating features reflecting low-temperature adaptation were effective but not enough to resolve the issue fully.

Next, we further investigated the factors causing inaccuracy in predicting the OGT of psychrophiles. We found that, while thermophiles were clustered in close clades (Blomberg’s K = 0.16), psychrophiles were relatively more sporadically distributed compared to thermophiles (Blomberg’s K = 0.04) in the phylogenetic tree (Figure 2a)^27, 28^. This result suggests that the OGT of psychrophiles has changed in a relatively short timescale. We estimated the evolutionary change in OGT of each species from their most recent common ancestor of the genus to which the focal species belongs (dOGT, see detail in Materials and Methods) (Figure 2b). All dOGT of psychrophiles were negative (mean dOGT = -11.0 °C), supporting the recent decrease in the OGT of psychrophiles (Figure 2c). It has been reported that the adaptation of amino acid composition requires the accumulation of numerous mutations and can take a long time^29, 30^. Therefore, we reasoned that rapid alterations in the OGT of psychrophiles may underlie the inaccuracy in the genome composition-based prediction. To test this hypothesis, we evaluated how rapid alterations in the OGT affect the prediction result in the next section.

**Figure 2.**
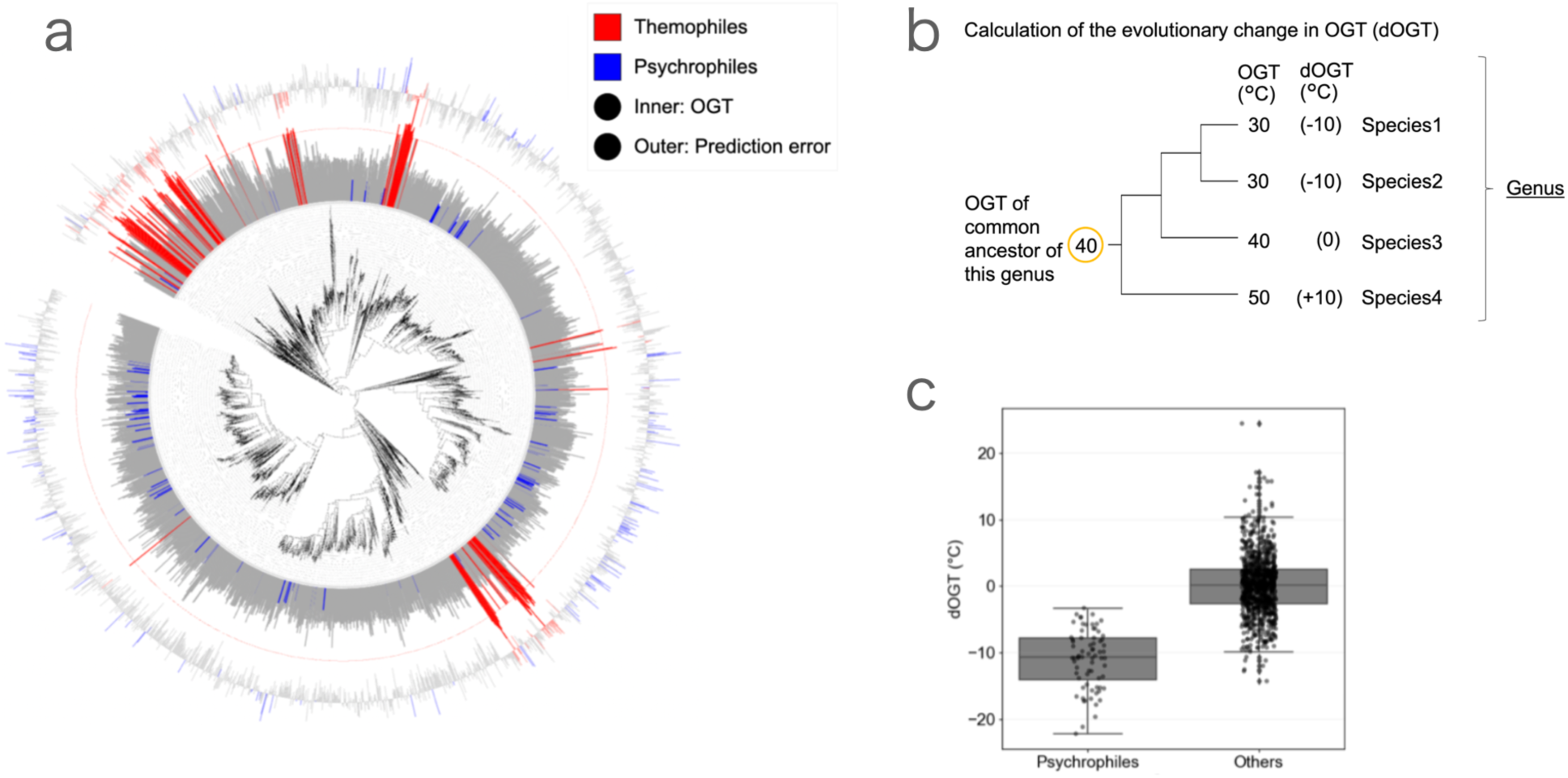
Calculation of the evolutionary change in OGT. **a** The distribution of OGT of bacteria with phylogenetic tree. The inner and outer bars indicate OGT and the prediction error of the model based on genome composition (Figure 1d), respectively. Red and blue bars indicate thermophiles (OGT > = 60 C°), and psychrophiles (OGT < 20 C°), respectively. The phylogenetic tree was obtained from GTDB r207. **b** The scheme of calculating the evolutionary change in OGT (dOGT) of each species. OGT of the common ancestor of each genus was calculated, and dOGT was defined as OGT_current–_OGT_ancestor._ **c** dOGT of psychrophiles (OGT < 20 C°) and other bacteria. Horizontal lines indicate median, boxes include second and third quartiles, and whiskers extend to points that lie within 1.5 times of the interquartile range. The dOGT of psychrophiles was –11.0 °C on average.

### The model based on genome composition cannot capture the diversity of OGT within the genus

We examined whether the model can predict the OGT of species that experienced rapid OGT alterations (Figure 3a,b). We found that the predicted OGT of species with large dOGT tends to be distinct from the observed OGT and that the predicted OGT is close to the ancestral OGT, even when OGT has increased or decreased from the ancestors (e.g., psychrophiles). For example, in *Marinobacter*, the observed OGT varies from 13 °C (*Marinobacter psychrophilus*) to over 45 °C (*Marinobacter lutaoensis*), while all predicted OGTs ranged from 25 °C to 39 °C ^31, 32^ (Figure 3c,d). In most genera, when comparing the species with the largest and lowest OGTs in the genus, the difference in predicted OGTs was small compared with the difference in the observed OGTs (Figure 3e).

**Figure 3.**
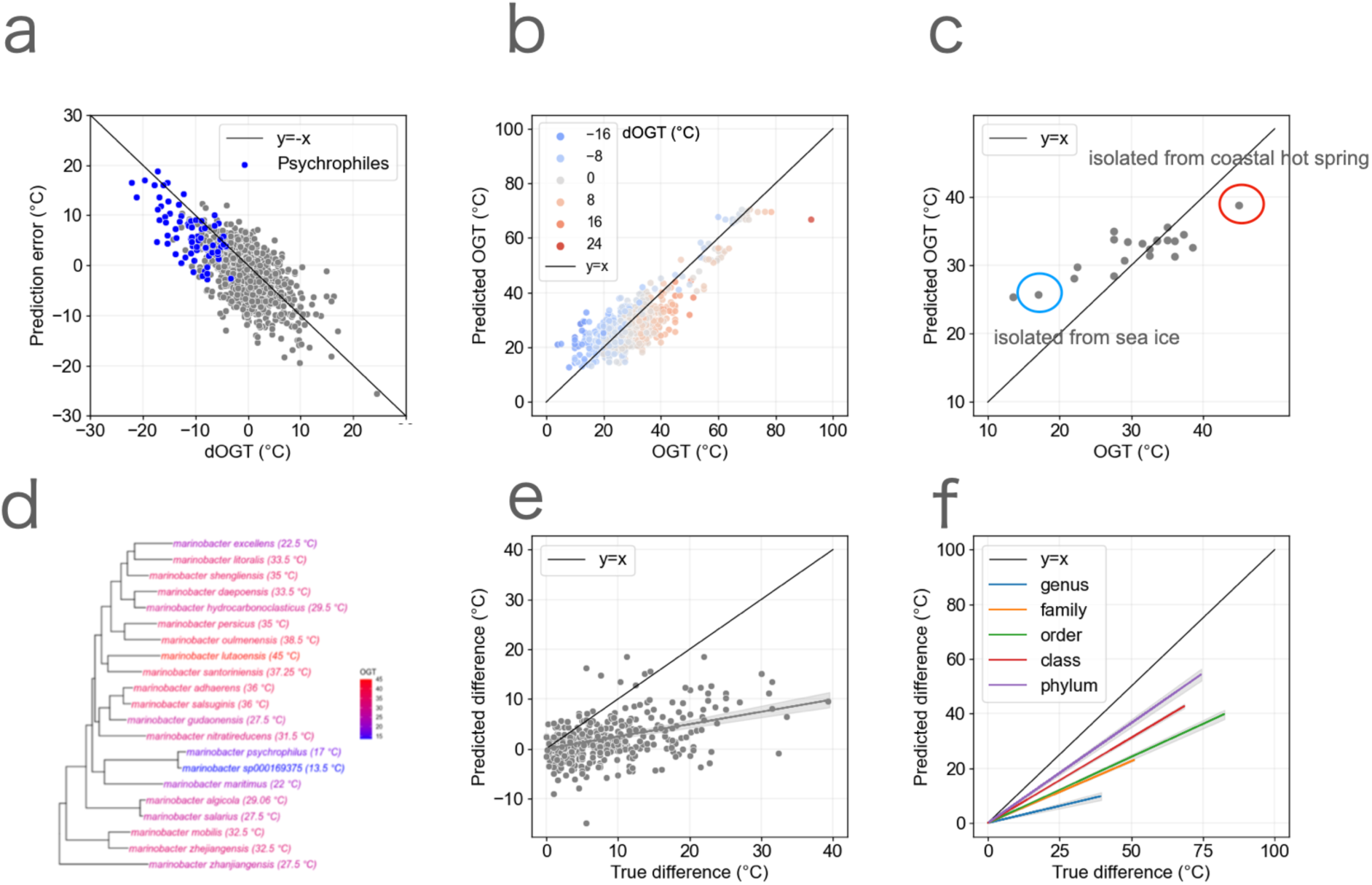
The model based on genome composition cannot fully capture the diversity of OGT within genera. **a** The prediction result (Figure 1d) is colored by dOGT. Blue and red indicate the species whose dOGT is under and over 0, respectively. **b** The relationship between dOGT and the prediction error (predicted OGT–observed OGT). Each point represents each species. The blue points represent psychrophiles. **c** The prediction of OGT for *Marinobacter*. *Marinobacter psychrophilus* (red circle) and *Marinobacter lutaoensis* (blue circle) were isolated from sea ice and coastal hot spring, respectively. **d** The phylogenetic tree obtained from GTDB r207, with OGT of the genus *Marinobacter.* The color represents OGT. **e** The relationship between the true OGT difference and the predicted OGT difference within the genus. Each point represents a genus. A pair of species that have the maximum and the minimum OGT in each genus was extracted, and the true OGT difference (the max OGT – the minimum OGT) and the predicted OGT difference (the predicted OGT of species with the maximum OGT – the predicted OGT of species with the minimum OGT) was calculated. The shade indicates the 90% confidence interval (CI) of the regression line. **f** The optimization of the genome composition over time. Pairs of two species were extracted within genus, family, order, class, and phylum. Similar to Figure 3e, the true difference of OGT and the predicted difference of OGT were calculated for each pair of species. The regression line was calculated separately for pairs within genus, family, order, class, and phylum. The shade indicates the 90% CI of the regression line.

If the genome composition adapts over time, the difference in predicted and observed OGTs should be mitigated as the phylogenetic distance between the two species increases. We extracted pairs of two species with various phylogenetic distances and compared the difference in predicted and observed OGTs, finding that the difference in predicted OGTs between two species became closer to the difference in the observed OGTs as the phylogenetic distance increased from genus, family, order, class, to phyla (Figure 3f). The inaccuracy of predicting the diversity of the OGT within the phylogenetically close clades may also suggest a potential bias in the machine-learning training process. To mitigate the potential bias, we conducted a similar analysis using the OGT prediction based on a simple regression of the raw values for each feature (MFE of tRNA, MFE of 16s rRNA, and amino acid composition of IVYWREL) (Supplementary Figure 3a, b, c). For all features, the difference in predicted OGTs became closer to the difference in the observed OGTs with increasing phylogenetic distance, suggesting that this result was not biased in the machine learning training process.

These results demonstrate that the inaccuracy in predicting the OGT of psychrophiles based on genome composition is attributed to the rapid decrease in their OGT, leading to insufficient optimization of genome composition.

### Using the information on the presence and absence of genes to improve the model

The genome composition of psychrophiles is often not optimized, making it difficult to predict their OGT based solely on genomic information. Nonetheless, psychrophiles exhibit lower OGT than their close relatives, suggesting that additional mechanisms contribute to their thermal adaptation. To explore this possibility, we focus on the presence or absence of genes associated with habitat temperature. Gene acquisition and loss are major factors in prokaryotic thermal adaptation^1, 9^, which has not been explicitly incorporated in previous studies^12, 14^. Thus, we tested whether integrating gene presence/absence information could improve prediction accuracy for psychrophiles.

Genes specific to a few clades may not be suitable for general predictions of OGT, particularly for uncultured bacterial clades whose OGTs remain poorly studied. Therefore, we focused on OGT-associated genes conserved across various bacterial lineages and OGT ranges. For this purpose, we selected species with relatively high and low OGT within each genus. We categorized them into a “high OGT group” (species with higher OGT within the genus) and a “low OGT group” (species with lower OGT within the genus). We then performed Fisher’s exact test to search for candidate genes associated with OGT (see Materials and Methods). Although genes broadly contributing to the change of OGT across various taxa were rare, K13993 (*Heat Shock Protein 20*), which is known to protect proteins against heat-induced denaturation and aggregation^33^, was significantly more often present in the high OGT group compared with the low OGT group (*p* = 4e-6 (0.04 after Bonferroni correction)) (Figure 4a, Supplementary Table 2). The phylogenetic analysis revealed that K13393 was widely distributed among bacterial lineages, suggesting its recurrent gains and losses (Figure 4b). To examine how the gains and losses of K13993 affect OGT, we inferred the ancestral state of the gene presence/absence. We compared it with the OGT of the ancestral species, finding that the loss of K13993 was associated with the decrease of OGT (-3.6 °C on average), although the acquisition of K13993 had little influence on OGT (-0.1 °C on average).

**Figure 4.**
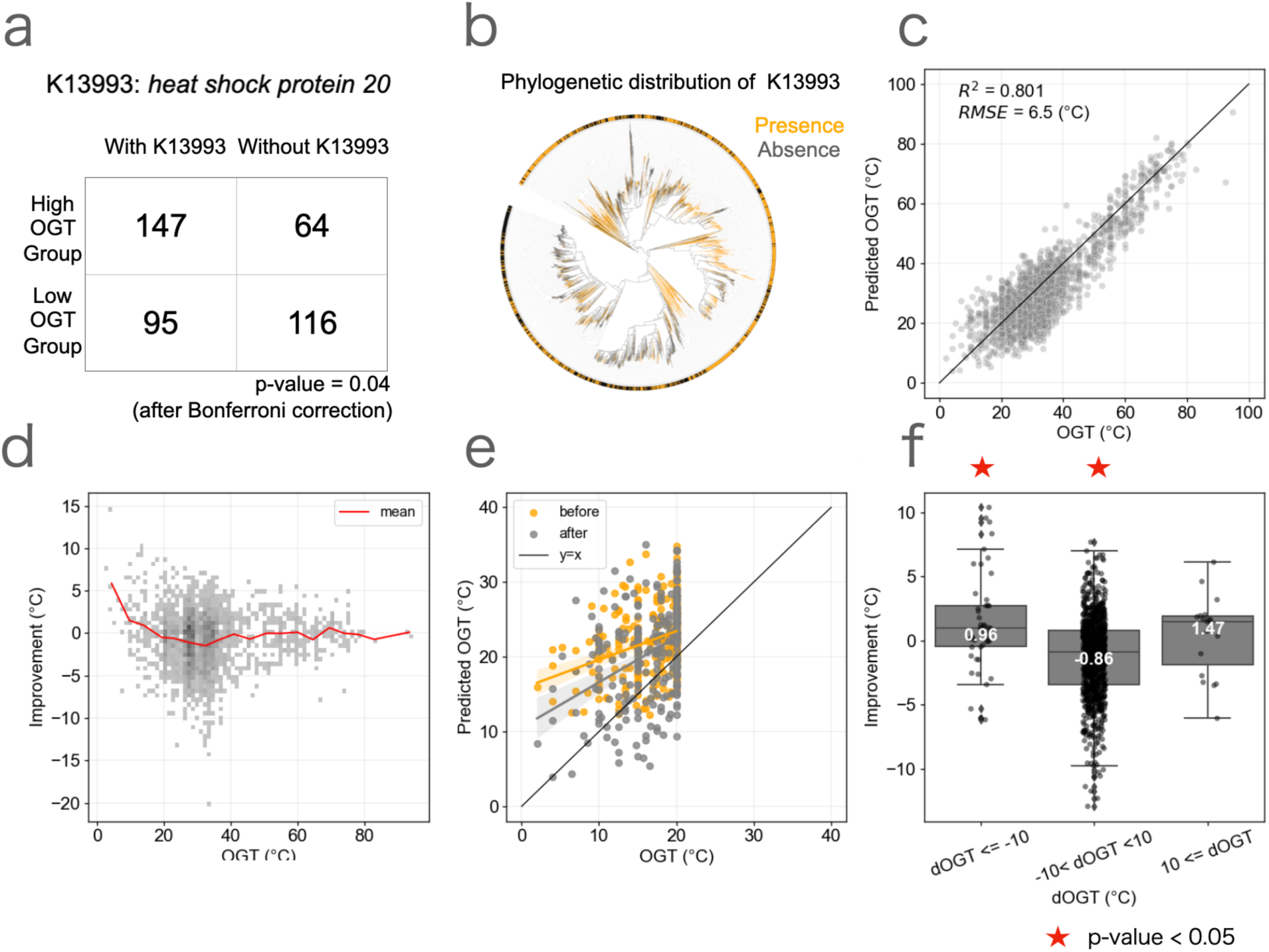
Inclusion of the gene presence/absence information improves the prediction accuracy. **a** A contingency table showing the association between the presence/absence of K13993 and OGT (*p* = 4.0 x 10^-6^ (0.04 after Bonferroni correction): Fisher’s exact test). **b** The distribution of K13993 in the phylogenetic tree obtained from GTDB r207. Orange and black indicate presence and absence of K13993, respectively. **c** The prediction by incorporating the presence and absence of the 100 genes into the model based on genome composition (Figure 1d). **d** The prediction improvement by incorporating the presence and absence of the 100 genes into the model based on genome composition (Figure 1d). The red line indicates the mean improvement of each OGT range. **e** The prediction improvement by incorporating the presence and absence of the 100 genes into the model based on genome composition (Figure 1d), particularly highlighting the improvements in psychrophiles. Orange and gray indicate before and after incorporating gene information, respectively. **f** The relationship between dOGT and the improvement of prediction after incorporating the presence/absence information of the 100 OGT-associated genes into the model based on genome composition (Figure 1d). Stars indicate that the mean values differed significantly from zero (*t*-test; *p* < 0.05). Horizontal lines indicate median, boxes include second and third quartiles, and whiskers extend to points that lie within 1.5 times of the interquartile range.

Although genes specific to cold adaptation are likely critical for improving the prediction of OGT in psychrophiles, such genes might not have been detected by the above method because of the limited number of psychrophiles in the dataset. To overcome this, we also searched for genes present in psychrophiles by comparing them with other bacteria (Table 1, Supplementary Table 3). The presence/absence of the K06970 (*23S rRNA (adenine1618-N6)-methyltransferase*) gene showed the strongest association (*p* = 5e-34 (5e-30 after Bonferroni correction)). This gene is known to help maintain the ribosomal peptide tunnel in the proper conformation^34^, suggesting that it can play an important role in structural stability at low temperatures. We confirmed the acquisition and the loss of K06970 was associated with the decrease (-2.5 °C on average), and the increase of OGT (2.7 °C on average), respectively. Furthermore, we found that the presence of two electron transport chain-related genes (K18159 and K08325) was also associated with cold temperature (Table 1). Those genes are suggested to associate the metabolic changes in low temperature environments^1, 35^. The pathway enrichment analysis of the top 50 genes revealed that psychrophiles tend to possess specific pathways related to anaerobic metabolism, such as methane and propionate metabolism (Supplementary Figure 4). In contrast, psychrophiles tended to lose genes in the aerobic processes such as pyruvate metabolism and the tricarboxylic acid cycle, possibly associated with the cold and anaerobic deep sea environments^36^. These results suggest that gene presence/absence information is a promising feature for capturing OGT changes that are not fully reflected in the genomic sequence optimization.

**Table 1.**
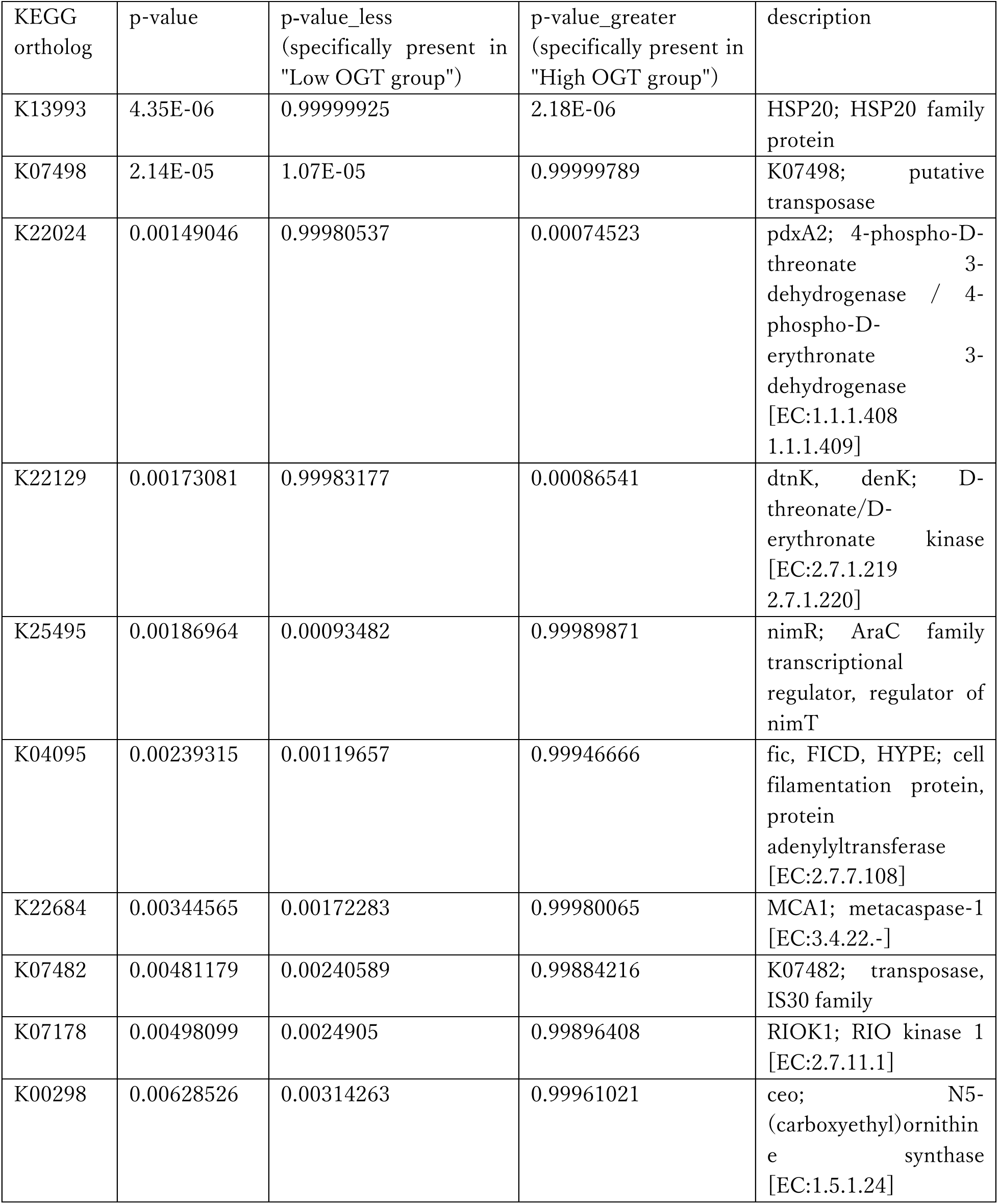
Top 10 candidate genes associated with changes in OGT across diverse lineages. KO numbers, p-values (with Bonferroni correction for multiple comparisons, significance level = 5.0E-6 (0.05/9959)), and descriptions for the top 10 genes are shown. Genes that are significantly present in either group were tested using Fisher’s exact test.

To evaluate whether incorporating gene presence/absence information improves the OGT prediction, we included the gene copy number information of the top 50 genes identified by each of the two methods described above (100 genes in total). For each training dataset, gene lists were generated independently to ensure no information was used from the phylum of the test data. By incorporating the gene copy number information, the accuracy in the prediction for the OGT of psychrophiles was improved (mean absolute error of psychrophiles = 6.3 °C) (Figure 4 c,d,e), particularly in the species with OGT < 5 °C (mean improvement = 7.6 °C). When compared with the prediction solely based on genome composition, the model demonstrated enhanced performance (1.1 °C on average) for species experiencing a rapid OGT shift (dOGT ≤ -10 °C and 10 °C ≤ dOGT), although the improvements of OGT of species with increase in OGT from ancestors (dOGT ≥ 10 °C) were not significant, which probably reflects the low number of species in this category (Figure 4f). Furthermore, the revised model better predicted the difference in OGT within genera and especially families compared with the previous model (Supplementary Figure 5). Conversely, accuracy did not improve for species whose dOGT was less than 5 °C (-0.86 °C on average, Figure 4f), resulting in a slight decrease in the overall accuracy (*R*^2^= 0.801). These results confirm that gene acquisition and loss significantly influenced the changes in OGT and demonstrate that incorporating gene presence/absence information improves the prediction of species experiencing the rapid OGT shift, especially psychrophiles.

### The conservation of OGT influences the correlation between OGT and genomic features

A previous study reported higher feature correlations with OGT in archaea than in bacteria, which makes archaea’s OGT more predictable^12^, partly because of the wide range of OGT in bacteria. However, the relationship of feature correlation with the conservation of OGT of bacteria and archaea has not been sufficiently explored. Because rapid alterations in OGT influence the accuracy of prediction as we demonstrated, the differences in the phylogenetic conservation of OGTs between archaea and bacteria would also be a key to determining the accuracy. Here, we compared OGT and genomic features (MFE of 16s rRNA, MFE of tRNA, and amino acids composition IVYWREL) in bacteria and archaea (Figure 5a). As previously reported^12^, the correlation between OGT and genomic features was stronger in archaea than in bacteria. To investigate the conservation of OGT in archaea, we mapped the OGT in archaea to the phylogenetic tree and compared the trait autocorrelation of bacteria and archaea (Figure 5b, c), finding that OGT was more conserved and stable in archaea than in bacteria.

**Figure 5.**
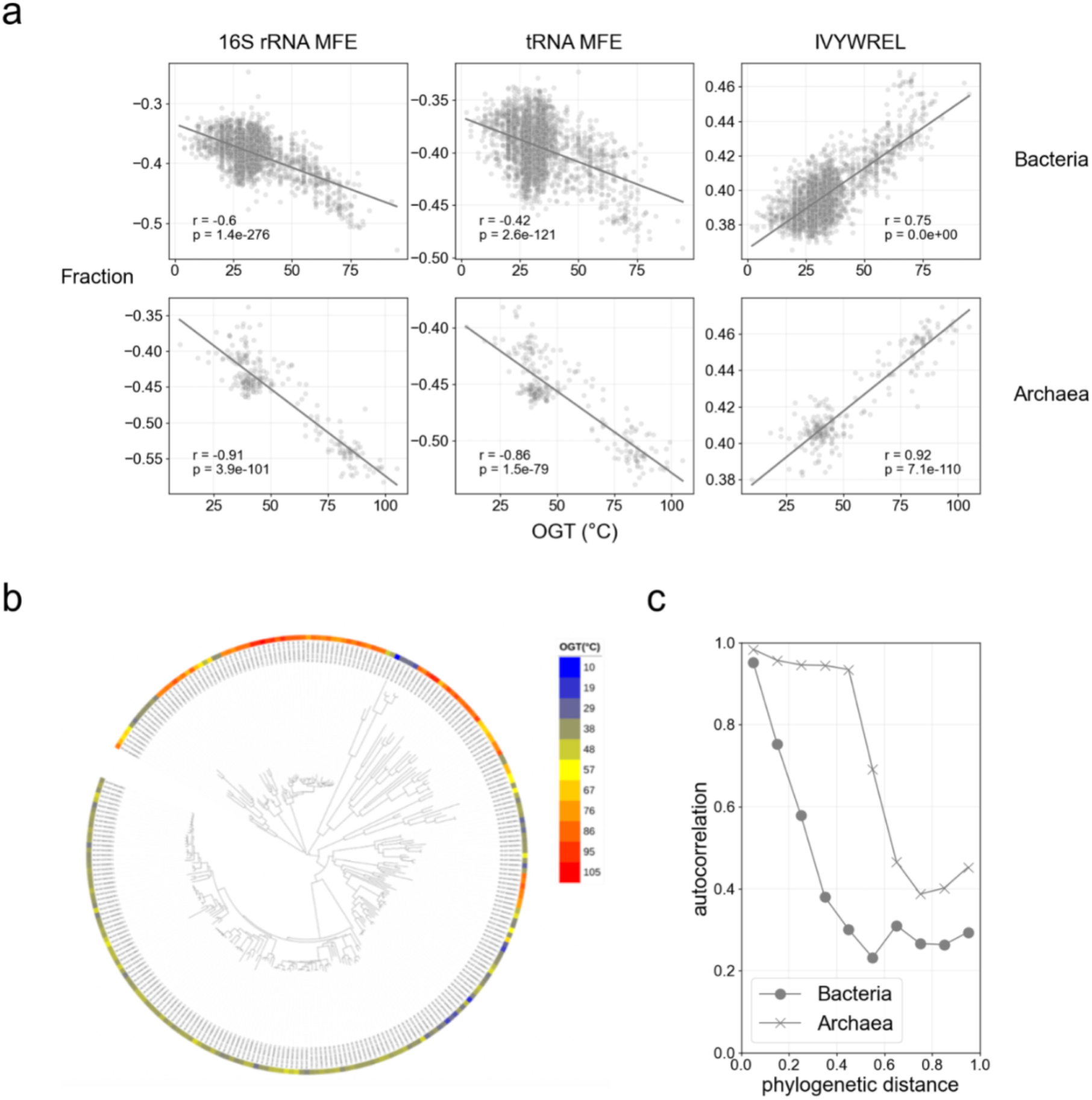
The conservation of OGT in bacteria and archaea. **a** The relationship between the genomic features (tRNA MFE, rRNA MFE, and summation of IVYWREL) and OGT. The top and bottom rows show bacteria and archaea, respectively. Pearson’s correlation coefficients and the p-values of the no-correlation test are shown. **b** The distribution of OGT of archaea in the phylogenetic tree obtained from GTDB r207. The color represents OGT. **c** The phylogenetic conservation of OGT of bacteria and archaea represented by autocorrelation in the tree. The circle and cross marks represent bacteria and archaea, respectively.

## Discussion

In this study, we demonstrated that incorporating gene presence/absence information into our model enhances the prediction of psychrophiles, which are often difficult to predict using genome composition alone^12^ (e.g., base composition, codon composition, and amino acid composition). Although genomic data from uncultivated prokaryotes have exponentially increased^11^, we do not know how many psychrophiles are hidden in those data. The ability to predict previously unknown psychrophiles may contribute to the discovery of new cold adaptation mechanisms and the identification of enzymes that remain active at low temperatures. Previous studies typically separated training and test data randomly, validating accuracy in this manner^14, 18^. However, since uncultured bacteria often belong to clades that are distinct from cultured bacteria^37^, it would be more appropriate to validate accuracy for clades not present in the training data. In this study, we employed leave-one-phylum-out cross-validation to ensure our model could predict the OGT of phyla absent from the training data. Interestingly, the linear model outperformed the non-linear model (SVR) under these conditions, indicating that the robustness of the linear model allowed it to capture genome composition features that are thermodynamically more meaningful. Those results suggest that our model may be applicable to rarely cultivated lineages such as candidate phyla radiation^38^.

The results suggest that the challenges in predicting psychrophiles primarily arise from the rapid decline in OGT from their ancestors. OGT can shift rapidly because of horizontal gene transfer or single nucleotide mutations^1, 39–42^, as laboratory evolution experiments have demonstrated^40, 41^. Contrary to this, changes in the genome composition often require the accumulation of numerous mutations and take considerable time^29, 30^. We propose that this difference in the mechanism of temperature adaptation causes the delay in the adaptation of genome composition to OGT, which influences the accuracy of predictions based on genome composition. Consistent with this idea, a recent study suggests that optimization of amino acid composition can also occur in psychrophiles over a long period when its lineage is ancestrally cold-adapted^30^. From this perspective, even though we identified features around the start codon reflecting cold adaptation (Figure 1c), these may not have undergone sufficient optimization. This could explain why incorporating features around start codons did not significantly enhance the prediction accuracy of psychrophiles. The results suggest that information on gene acquisition and loss is valuable for predicting OGT diversity within genera, leading to the improvement of the prediction of OGT of psychrophiles. While some of the genes whose presence/absence was associated with OGT could be directly involved in temperature adaptation, many of them are functionally uncharacterized, which require further experimental investigation. We note that not only presence/absence but also single nucleotide substitutions can contribute to the changes in OGT^40, 41^, and future models could incorporate their effects into the predictions, potentially improving accuracy. While we focused on OGT in this study, similar challenges may arise in predicting rapidly altered phenotypes for other traits, such as optimal pH and salinity, as these are also known to be influenced by the gain and loss of specific genes^29, 43^. The present study suggests that gene information may also be important for predicting such phenotypes.

Previous studies discussed that the OGT of archaea is more accurately predictable than those of bacteria^12^. Although this phenomenon can be caused by multiple factors such as the range of OGT ^12, 44^, our finding of the high conservation of OGT in archaea suggests that their genomes have undergone long-term optimization. A previous study suggested that evolution in archaea tends to be relatively stable because of their inherent advantages in adapting to chronic energy stress, whereas bacterial evolution is more dynamic^45^.

In our study, we proposed that a model based on genome composition poses a challenge in predicting the OGT of species whose OGT has rapidly changed, and that combining gene presence/absence information would be the key to predicting the OGT of psychrophiles. Although machine-learning models based on genome composition for phenotype prediction have gained attention in recent years^12, 14, 46^, the present study highlights the importance of incorporating gene sets related to phenotype into predictions.

## Materials and Methods

### Dataset of OGT and genome

Bacterial and archaeal OGT data were obtained from three publicly available datasets (I: 11,004 species ^18^, II: 8,639 species ^16^, and III: 4,663 species ^17^). All data were downloaded on October 3, 2022. Most of these OGT data are experimentally determined optimal incubation temperatures. The species with OGTs exactly matching 25 °C, 28 °C, 30 °C, and 37 °C were most frequently observed in all three databases. We excluded those species from the analysis because these temperatures are often used as conventional incubation temperatures in laboratories and seem to be an artifact caused by hand-built bias^47^. As a result, the number of species was 4,038 for the dataset I, 4,312 for II, and 2,527 for III. The three databases were merged, and overlapping species were averaged. The merged data resulted in 8,972 species. The OGT data were merged with representative genomes from the Genome Taxonomy Database (GTDB) r207^19–22^ with completeness > 95 % and contamination < 5 % using the species name as a key because OGT data with genome accession were only available for 7 % of species (637 species). Using OGT data of the species with more than three strains^17^, we ensured the diversity of OGT within species was relatively small (median of standard deviation = 0.52 °C).

Coding sequence annotations were downloaded from GTDB r207 with data predicted by Prodigal 2.6.3^48^. tRNA annotation was performed using tRNAscan-SE 2.0.9.^49^ and rRNA annotation was performed using barrnap 0.9 (https://github.com/tseemann/barrnap). Excluding species for which some of the 5S rRNA, 16S rRNA, and 23S rRNA annotations were missing, the data of 2,869 bacterial and 262 archaeal species were used in this study. The taxonomy of each species was defined based on GTDB taxonomy. The phylogenetic tree was obtained from GTDB r207 and was drawn using iTOL (interactive Tree of Life) v6^50^. For rRNA and tRNA, the most stable structures and minimum free energies were obtained using Linearfold^51^ (parameters were set to -V and a Vienna RNAfold-based model was used).

### Downsampling of the mesophiles

The distribution of bacterial OGT is highly biased toward the mid-temperature range (20 °C to 40 °C). Therefore, we also made the downsampled dataset of bacteria (Figure 1a). For the downsampling, the species with the highest genome quality score (completeness - 5 × contamination) were sampled from each genus for species with OGT between 20 °C and 40 °C. With this downsampling, 1,631 species were kept, and training and evaluation of regressions were conducted using this dataset. To avoid the influence on the biological interpretation, the prediction result of all data was used for the following analysis.

### Extraction of specific information near start codons

The following method was used to extract site-specific changes near the start codon. *F_i_* is the features of the average base composition (*i*-th to *i*+9-th) and amino acid composition (*i*-th) counting from the start codon. The mean value of all genes was calculated for each species. To extract the site-specific information of *F_i_*, we normalized *F_i_* using the representative value of *F_rep_*:

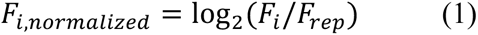

The *F_rep_* value was the 50th value from the start codon when analyzing base composition, and the average of total length sequence when analyzing amino acid composition. We used *F*_5._ as the representative feature for guanine composition near the start codon, and the mean of *F*_1_ - *F*_4_ as the representative feature for amino acid composition near the start codon.

### Creating regression models based on genome composition

We calculated a total of 526 features from the genomes (Table 1). All features were normalized to a maximum value of 1 and a minimum value of 0. For machine learning, Lasso regression and SVR were used (Scikit-learn 1.2.2). The parameter with the best score with features was adopted (Lasso: alpha = {0.002, 0.004, 0.006, 0.008}, SVR: C = {0.1, 1, 10, 100}). To verify the accuracy for unknown clades, phylum-out cross-validation was used, in which one phylum was the test data, and the rest were the training data, and validation was performed so that all phyla were the test data one by one. Root mean squared error (RMSE) and coefficient of determination (*R*^2^) were used to evaluate the prediction accuracy.

### Ancestral state reconstruction of OGT and genes

The OGT of most recent common ancestor of each genus was estimated by minimizing the summation of squared changes in OGT along the branches of the tree (“asr_squared_change_parsimony()” function from R package castor 1.7.11 with the parameter “weighted” set to True). By calculating the difference between the OGT of the common ancestor of each genus (with more than five species) and the OGT of the current species (*OGT_current_species_* − *OGT_common_ancestor_*), we calculated how much the OGT of that species has changed from the ancestor (dOGT). Similarly, the ancestral state of genes (presence or absence) was inferred using the function “asr_mk_model ()” function. The trait autocorrelation of OGT was calculated using the “get_trait_acf()” function.

### Comparing the true OGTs and predicted OGTs in phylogenetically close clades

To verify how accurately the predicted OGT captured the changes in OGT of closely related species, we compared two species with the highest and lowest OGTs in the genus, and generated the scatter plot of the difference in true OGTs and predicted OGTs. We calculated the 90 % confidence interval of the slope by a linear regression with intercept = 0 and bootstrap sampling of 100 times. Next, we compared the two species that belong to the same family but different genera and have the largest difference in true OGT. Those species pairs were taken for all families and all combinations of genera that belong to each family. The method described above was applied similarly for the two species that belong to the same order, the same class, the same phylum, and the same domain.

### Annotation of gene and gene set enrichment analysis

The CDSs of the genome were functionally annotated using KofamScan version 1.3.0 with default parameters^52^. For the best functional prediction, the default asterisk annotation was adopted. To search for genes related to OGT, we focused on genera that included both high and low OGT species. Specifically, pairs of species with the highest and lowest OGT in a focal genus were extracted iteratively until the difference was less than 10 degrees, with a maximum of three pairs per genus to alleviate phylogenetic bias. We then performed Fisher’s exact tests with Bonferroni corrections to assess the statistical significance of the association between OGT and the gene presence/absence. Similarly, we performed Fisher’s exact tests for psychrophiles (OGT < 20°C) and other bacteria under the same criteria. KEGG pathway enrichment analysis was conducted using R package clusterProfiler version 4.14.4^53^.

## Acknowledgments

This research was supported by JST DICP Grant Number. JPMJND2206 and GteX Grant Number JPMJGX23B2 to MM and JSPS KAKENHI Grant Numbers JP22H04925 (PAGS) to WI., 19H05679, 19H05688, 23KK0122, and 24K01889 to MM, 22K21352 and 23K27228 to TT. Computational resources were partially provided by the NIG supercomputer at the ROIS National Institute of Genetics.

## Notes

### Competing Interest Statement

The authors have declared no competing interest.

