## Supplementary Figures 1-5 for "Genomic and evolutionary factors influencing the prediction accuracy of optimal growth temperature in prokaryotes"

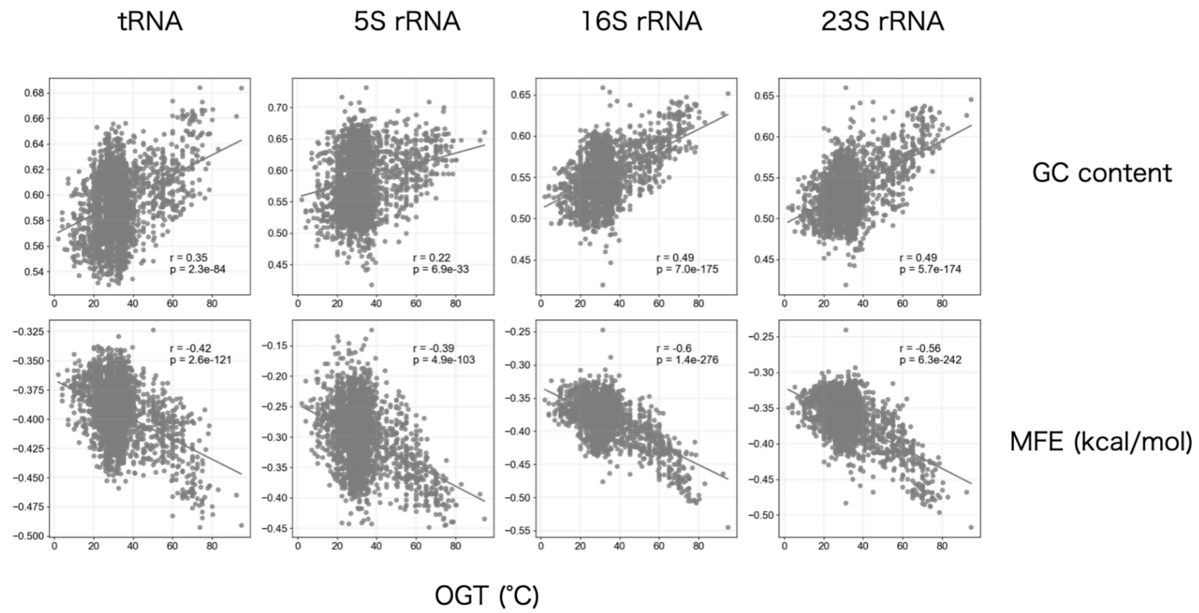


**Supplementary Figure 1. The relationship between OGT, GC content, and MFE of tRNA and rRNA.** MFE of tRNA and rRNA were normalized by sequence length. Pearson's correlation coefficients and the p-values of the no-correlation test are shown.

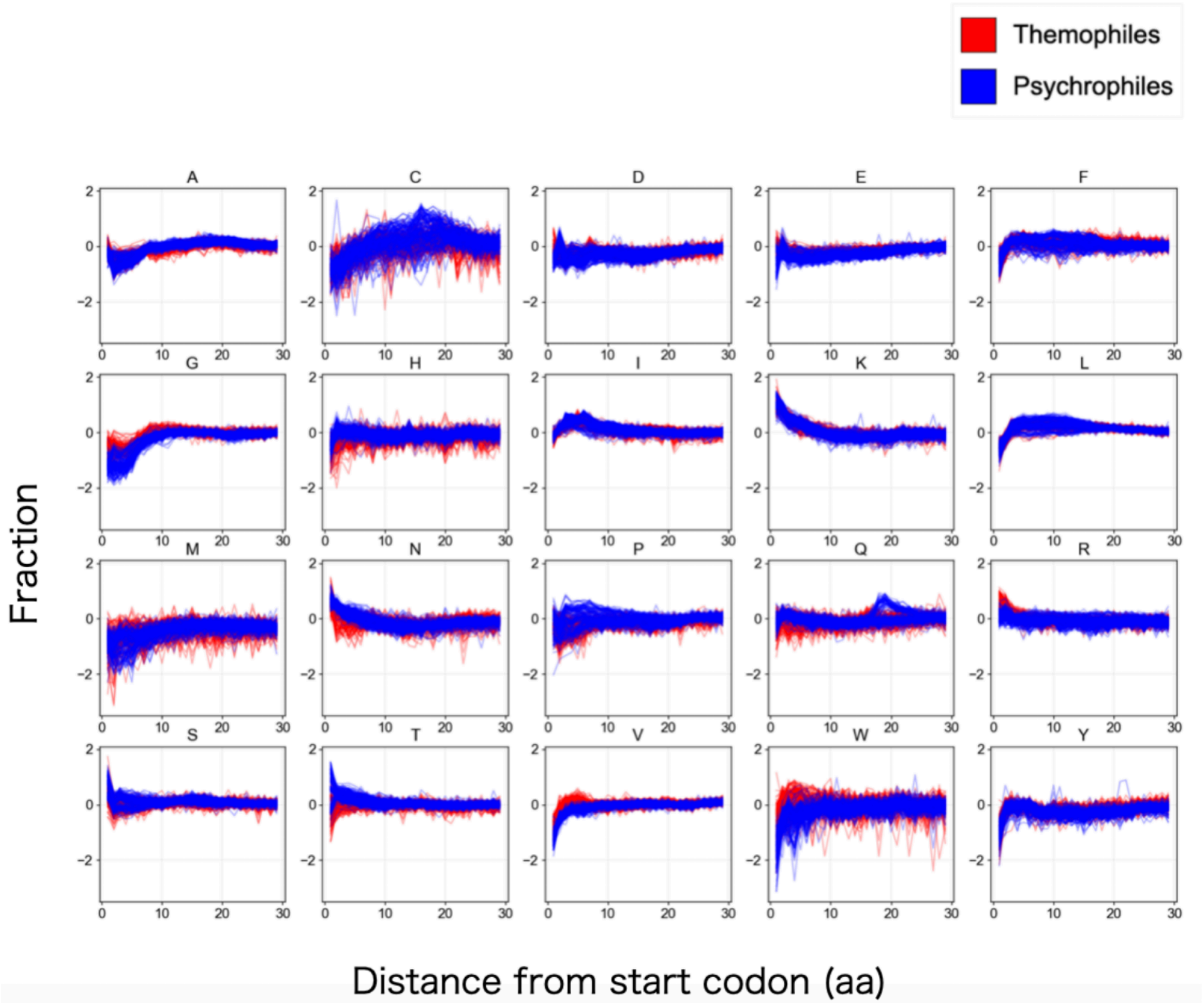


**Supplementary Figure 2. The fraction of amino acids in the downstream region of the start codon.** The fraction was normalized by the average of the fraction of each amino acid of species. The red and blue lines indicate thermophile and psychrophile, respectively.

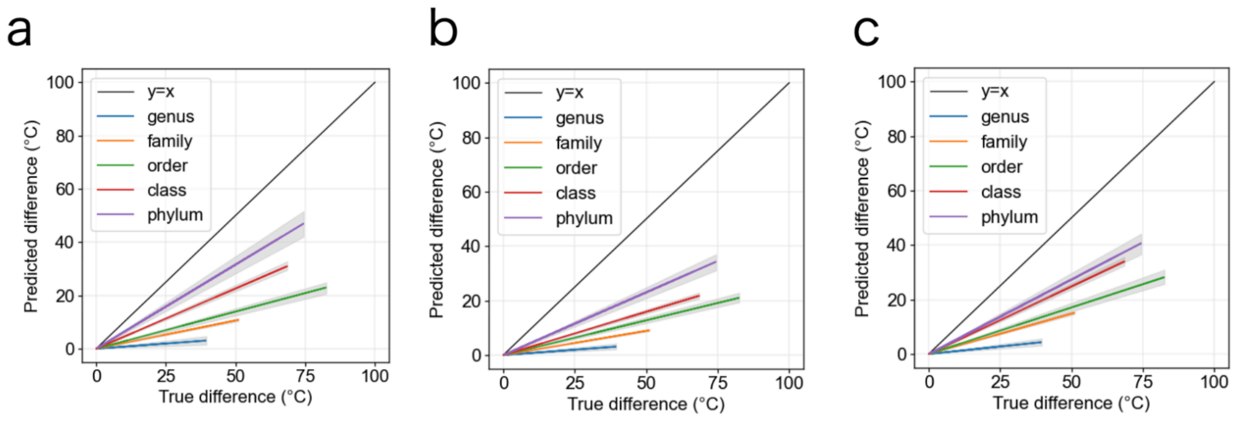


**Supplementary Figure 3. The optimization of the MFE of 16S rRNA, MFE of tRNA, and summation of IVYWREL over time.** Each figure was generated with the prediction result by simple regression using MFE of 16S rRNA (**a**), MFE of tRNA (**b**), and summation of IVYWREL (**c**). To avoid overfitting to mesophilic temperatures, species with mesophilic OGT (20–40°C) were excluded during training.

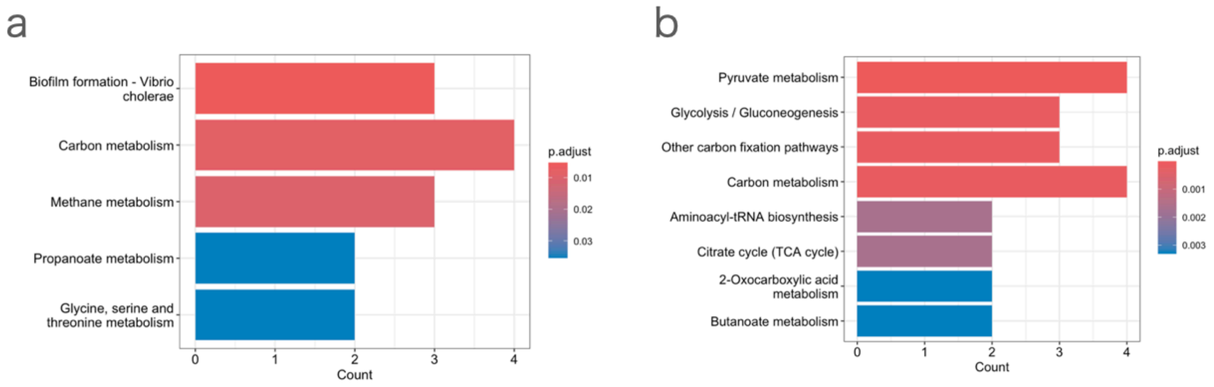


**Supplementary Figure 4. The pathway enrichment analysis of the top 50 cold adaptation-associated genes (adjusted p-value < 0.01).** Enrichment pathways with genes that are specifically present (**a**) or absent (**b**) in psychrophilces.

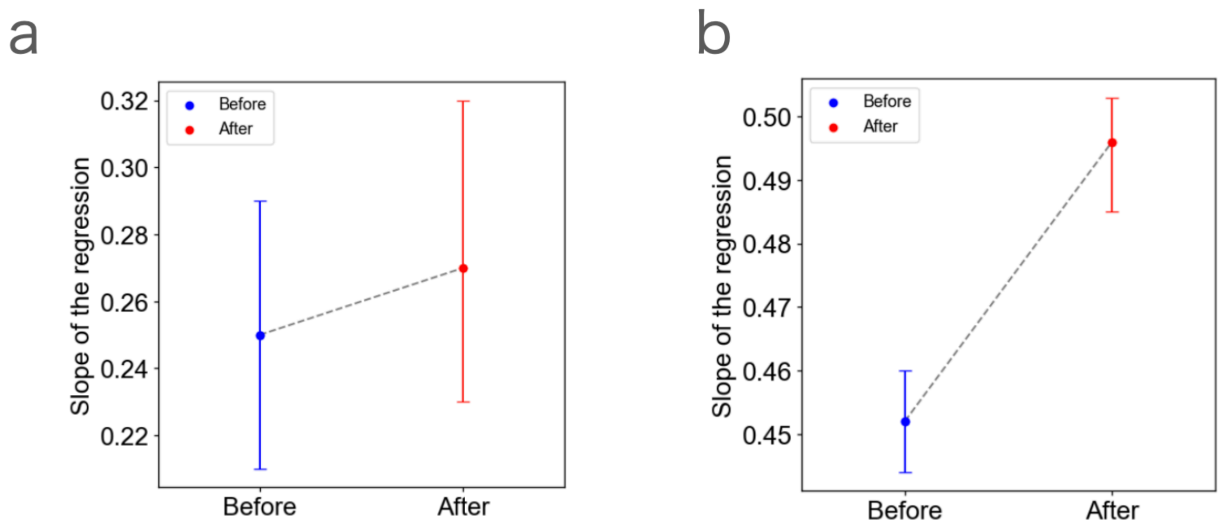


**Supplementary Figure 5. The improvement of the prediction of OGT with the information on the presence and absence of OGT-associated genes.** The slope of the regression for the true OGT difference and the predicted OGT difference within each genus (**a**) and family (**b**) (same method as the slope of the regression line of the genus and the family in Figure 3f) before and after incorporating OGT-associated gene information with a 90% confidence interval.
